# The molecular landscape of pediatric acute myeloid leukemia reveals recurrent structural alterations and age-specific mutational interactions

**DOI:** 10.1101/125609

**Authors:** Hamid Bolouri, Jason E. Farrar, Timothy Triche, Rhonda E. Ries, Emilia L. Lim, Todd A. Alonzo, Yussanne Ma, Richard Moore, Andrew J. Mungall, Marco A. Marra, Jinghui Zhang, Xiaotu Ma, Yu Liu, Yanling Liu, Jaime M. Guidry Auvil, Tanja M. Davidsen, Patee Gesuwan, Leandro C. Hermida, Bodour Salhia, Stephen Capone, Giridharan Ramsingh, Christian Michel Zwaan, Sanne Noort, Stephen R. Piccolo, E. Anders Kolb, Alan S. Gamis, Malcolm A. Smith, Daniela S. Gerhard, Soheil Meshinchi

**Affiliations:** Division of Human Biology, Fred Hutchinson Cancer Research Center, Seattle, WA; Winthrop P Rockefeller Cancer Institute, University of Arkansas for Medical Sciences and Arkansas Children’s Research Institute, Little Rock, AR; Jane Anne Nohl Division of Hematology, USC/Norris Comprehensive Cancer Center, Los Angeles, CA; Clinical Research Division, Fred Hutchinson Cancer Research Center, Seattle, WA; Canada’s Michael Smith Genome Sciences Centre, British Columbia Cancer Agency, Vancouver, BC, Canada; Keck School of Medicine, University of Southern California, Los Angeles, CA; Children’s Oncology Group, Monrovia, CA; Division of Computational Biology, St Jude Children’s Research Hospital, Memphis, TN; Office of Cancer Genomics, National Cancer Institute, Bethesda, MD; Department of Translational Genomics, Keck School of Medicine, University of Southern California, Los Angeles, CA; Dept of Pediatric Oncology, Erasmus MC-Sophia Children’s Hospital, Rotterdam; Department of Biology, Brigham Young University, Provo, UT; Department of Biomedical Informatics, University of Utah, Salt Lake City, UT; Nemours Center for Cancer and Blood Disorders, Alfred I. DuPont Hospital for Children, Wilmington, DE; Division of Hematology/Oncology/Bone Marrow Transplantation, Children’s Mercy Hospitals and Clinics, Kansas City, MO; Cancer Therapy Evaluation Program, National Cancer Institute, Bethesda, MD

## Abstract

We present the molecular landscape of pediatric acute myeloid leukemia (AML), characterizing nearly 1,000 participants in Children’s Oncology Group (COG) AML trials. The COG/NCI TARGET AML initiative assessed cases by whole-genome, targeted DNA, mRNA, miRNA sequencing and CpG methylation profiling. Validated DNA variants revealed diverse, infrequent mutations with fewer than 40 genes mutated in >2% of cases. In contrast, somatic structural variants, including novel gene fusions and focal *MBNL1*, *ZEB2*, and *ELF1* deletions, were disproportionately prevalent in young as compared to adult patients. Conversely, *DNMT3A* and *TP53* mutations, common in adults, are conspicuously absent from virtually all pediatric cases. Novel *GATA2*, *FLT3*, and *CBL* mutations, recurrent *MYC-ITD, NRAS, KRAS*, and *WT1* mutations are frequent in pediatric AML. Deletions, mutations, and promoter DNA hypermethylation convergently impact Wnt signaling, Polycomb repression, innate immune cell interactions, and a cluster of zinc finger genes associated with *KMT2A* rearrangements. These results highlight the need for, and facilitate the development of age-tailored targeted therapies for the treatment of pediatric AML.

---

Acute leukemia is the most common form of childhood cancer^1^, and its incidence is increasing. Despite constituting only 20% of pediatric acute leukemia, acute myeloid leukemia (AML) is overtaking acute lymphoblastic leukemia (ALL) as the leading cause of childhood leukemic mortality, in part because current prognostic schemas classify many children who will ultimately succumb to their disease as low-or intermediate-risk. Additionally, aside from investigational tyrosine kinase inhibitors for *FLT3*-activated AML, targeted therapies are not used in pediatric AML. Both problems stem from an inadequate understanding of the biology of childhood AML.

AML is a molecularly heterogeneous group of diseases affecting patients of all ages^2^. Recent genome-scale studies have revealed novel, potentially targetable mutations prevalent in adult *de novo* AML^3–5^. However, the relevance of these findings to childhood AML remains unclear, since several of the most common adult mutations appear far less prevalent in pediatric AML^6,7^.

To date, no comprehensive characterization of pediatric AML has been described. Here, we report the initial results of the TARGET (Therapeutically Applicable Research to Generate Effective Treatments) AML initiative, a collaborative COG/NCI project to comprehensively characterize the mutational, transcriptional, and epigenetic landscapes of a large, well-annotated cohort of pediatric AML. Comparing AML molecular profiles across age groups, we show that stark differences in mutations,d structural variants and DNA methylation distinguish AML in infants, children, adolescents, and adults.

## Results

### Overview of cohort characteristics

A total of 1023 children enrolled in COG studies are included in the TARGET AML dataset. Comprehensive clinical data, including clinical outcomes and test results for common sequence aberrations (outlined in **Table S1**), are available for 993 patients. Of these, 815 subjects were profiled for somatic mutations at presentation: 197 by whole-genome sequencing (WGS), and 800 by targeted capture sequencing (TCS), at read depths averaging 500x, for validation of mutations identified by WGS. The WGS discovery cohort of diagnostic and remission (germline comparison) specimens were selected from patients treated on recent COG studies who achieved an initial remission to induction chemotherapy. These trials randomized type or timing of induction therapy (CCG-2961)^8^ and the addition of gemtuzumab ozogamicin in a single arm pilot (AAML03P1)^9^ or randomized fashion (AAML0531)^10^. Specimens for TCS validation were obtained from 800 patients, including 182 from the WGS discovery cohort (153 with matched remission samples). A complete listing of cases and their characterization is available in the TARGET sample matrix (https://ocg.cancer.gov/programs/target/data-matrix). The age at presentation of TARGET AML participants ranged from 8 days to 29 years (median 10 years, Fig. 1a). Infants (<3 years old), children (age 3-14) and adolescents/young adults (AYA; age 15-39) differ broadly by cytogenetic and clinical risk-group classifications (Fig. 1a, multivariate Chi-squared p<10^−22^), consistent with observed differences in clinically-evaluated structural abnormalities and mutations (summarized in Fig. 1b). Notably, among these clinically detected abnormalities, only 5 mutations and 5 structural aberrations occur in more than 5% of patients (mutations in *FLT3, NPM1, WT1, CEBPA*, and *KIT*; fusions involving *RUNX1, CBFB* and *KMT2A*; trisomy 8 and loss of the Y chromosome.)

**Figure 1.**
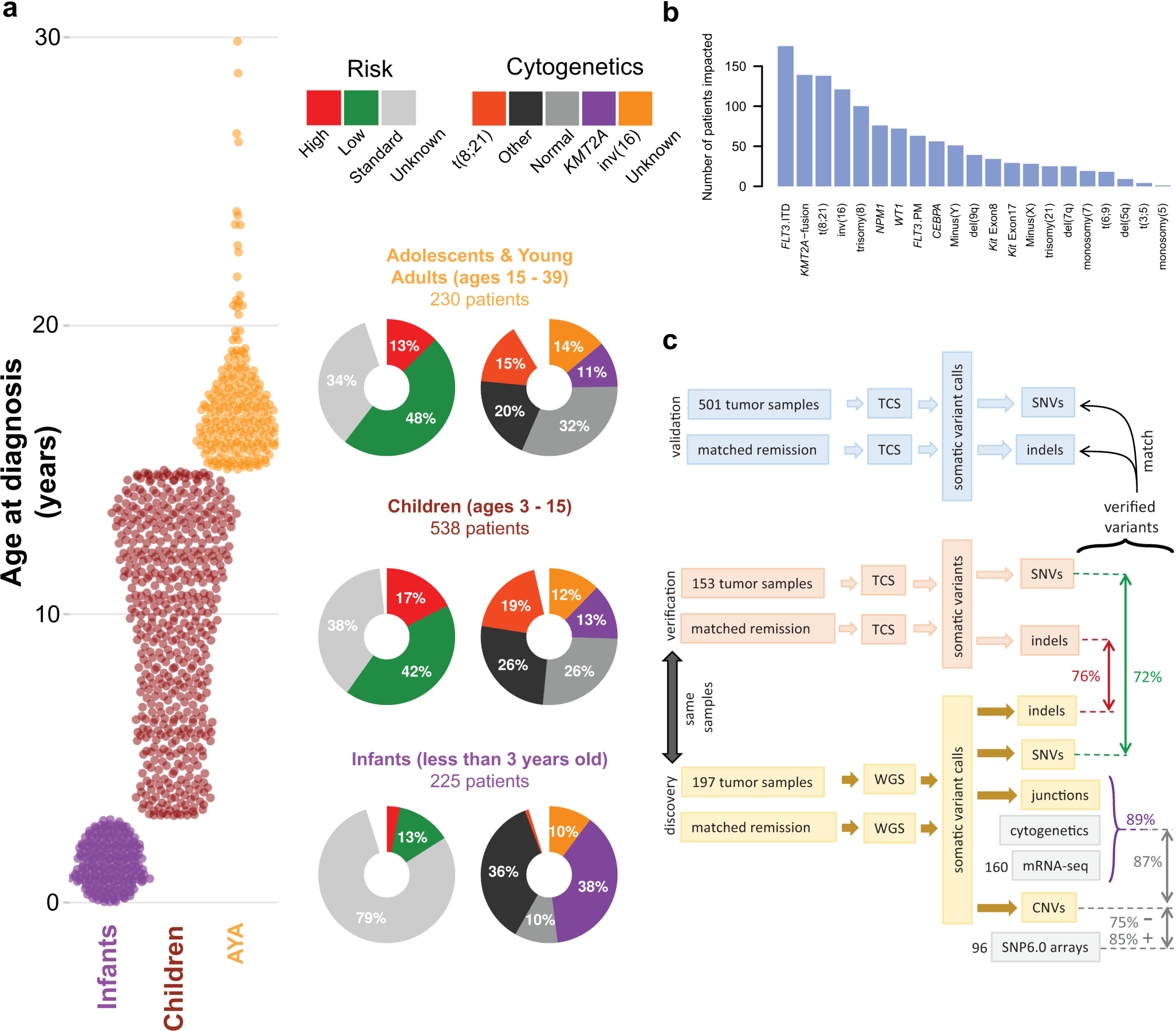
An overview of the TARGET AML study. (**a**) The distribution of subjects by clinical risk category and cytogenetic classification is shown adjacent to each age group analyzed (Infant, <3 years; Child, 3 to <15 years; Adolescent/Young Adult (AYA), 15 to <40 years). (**b**) A summary of the clinically established molecular aberrations in the cohort (n=993) is illustrated. FLT3.ITD, *FLT3* internal tandem duplications, FLT3.PM, *FLT3* D835 point mutations. (**c**) Overview of the genomic variant discovery, verification, and validation process. We characterized diagnostic and remission (taken as germline) samples from 197 patients using whole genome sequencing (WGS) and verified 153 diagnostic/remission case pairs by targeted capture sequencing (TCS) of genes recurrently impacted in the WGS samples (an additional 29 WGS cases were verified by TCS of diagnostic cases only, see **Fig. S1**). 72% of WGS SNVs, and 76% of WGS indels were confirmed by TCS (red & green text in figures). For focal copy number (CN) alterations spanning fewer than 7 genes, 75% of recurrent WGS deletion/loss and 85% gain/amplification calls matched recurrent alterations discovered by SNP6 arrays in 96 matching samples. For chromosomal junctions, we integrated WGS, clinical karyotyping and RNA-seq data by majority vote, confirming 89% of WGS junction calls.

We validated each class of somatic DNA sequence alteration discovered by WGS through secondary assays (Figs. 1c and **S1**). Single nucleotide variants (SNVs) and short insertions and deletions (indels) were confirmed by TCS of the coding sequences of the genes identified as recurrently altered in the WGS studies. WGS-detected copy number alterations were confirmed by GISTIC2.0 scores from SNP arrays; WGS-detected structural changes (such as translocations and inversions) were confirmed by RNA-seq and clinical leukemia karyotyping data. Across variant types, we find >70% concordance between at least two assays. These variants are referred to as verified variants hereon. An overview of the multiplatform-verified somatic DNA variants in 684 patients is presented in Fig. 2a. Roughly a quarter of patients possess normal karyotype, yet nearly all revealed at least one recurrent verified somatic DNA alteration, and at least 12 common cancer-associated cellular processes are recurrently impacted (**Fig. S2**, **Tables S2a**, **b**).

**Figure 2.**
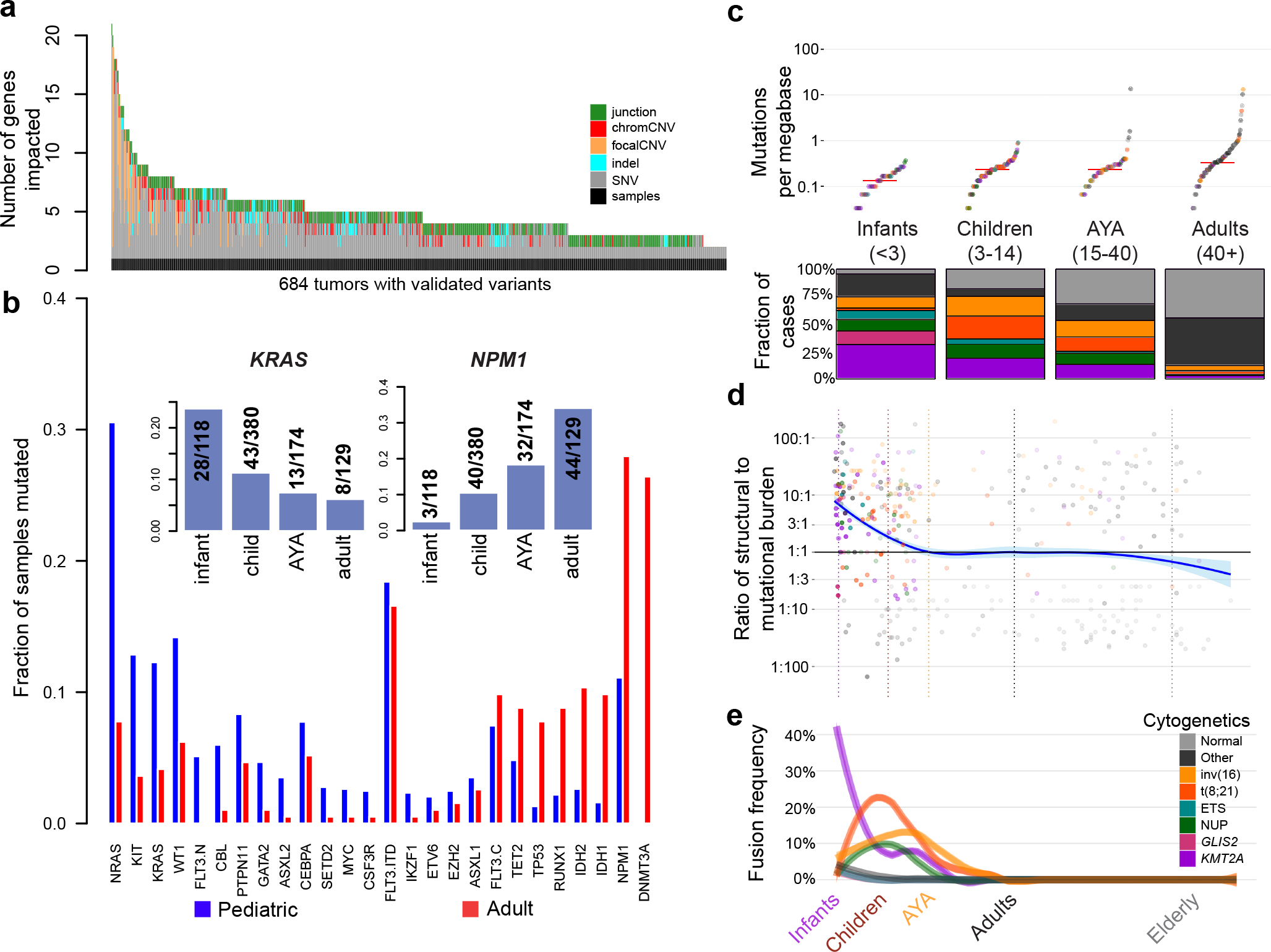
Age-related differences in mutational and structural alterations in AML. (**a**) Distribution of variants per sample. At least one variant impacting a gene recurrently altered in pediatric AML was identified by multi-platform validated variants in 684 patients. Junction, protein fusions (see methods); chromCNV, chromosomal arm/band level copy variant; focalCNV, gene level copy variant; indel, small insertion/deletion; SNV, single nucleotide variant. (**b**) Age-dependent differences in the prevalence of mutations. *FLT3* mutations are plotted in 3 categories: internal tandem duplication (ITD; FLT3.ITD), activation loop domain (FLT3.C), and novel, childhood-specific changes (FLT3.N). (**b, inset**) A pattern of waxing or waning mutation rates across age groups is evident in selected genes (*KRAS* and *NPM1* illustrated). (**c**) Childhood AML, like adult AML, has a low somatic mutational burden (top and **Fig. S5**), but is more frequently impacted by common cytogenetic alterations (lower section). For color key, see legend at bottom-right. (**d**) The ratio of the burden of structural variation to SNVs/indels is high in infancy and early childhood and declines with age. For color key, see legend at bottom-right. (**e**) Using a sliding-window approach to account for uneven sampling by age, the incidence of common translocations in AML is shown to follow age-specific patterns (multi-variate Chi-squared p < 10^-30^), and to be greatest in infants compared to all other ages (Chi-squared p < 10^−22^). *KMT2A* fusions are most common in infants (Chi-squared p < 10^−20^), while core binding factor fusions tend to affect older children (Chi-squared p < 10^−7^).

We carried out analyses of microRNA, mRNA, and/or DNA methylation in 412 subjects. A summary of the assays performed and case-assay overlap is presented in **Fig. S3**. We compared our verified variants to those of 177 adult AML cases from The Cancer Genome Atlas (TCGA) project^3^, stratified by the age groupings outlined in Fig. 1a. The TARGET and TCGA discovery cohorts both contained numerous AYA patients (**Table S3**). Importantly, our conclusions regarding the molecular characteristics of this age group are identical when analyzing either or both cohorts (**Fig. S4**).

### Somatic gene mutations in pediatric AML

Like adult AML, pediatric AML has one of the lowest rates of mutation among molecularly well-characterized cancers (**Fig. S5**), with < 1 somatic, protein-coding change per megabase in most cases. However, the landscape of somatic variants in pediatric AML is markedly different from that reported in adults^3,4^ (Figs. 2b, **S6-S7**, **Table S4**). *RAS, KIT,* and *FLT3* alterations, including novel, pediatric-specific *FLT3* mutations (FLT3.N), are more common in children. Mutational burden increases with age, yet older patients have relatively fewer recurrent cytogenetic alterations. Indeed, the number of coding SNVs, within and across cohorts, is best predicted by age (Fig. 2c, p<10^−15^) and by cytogenetic subgroup. In contradistinction to the higher prevalence of small sequence variants in older patients, recurrent structural alterations, fusions, and focal copy number aberrations are more common in younger patients (Figs. 2d-e, p<10^−3^, see below). Patients with *CBFA2T3-GLIS2, KMT2A,* or *NUP98* fusions tend to have fewer mutations (p<10^−9^), with subgroups demonstrating inferior clinical outcome (**Fig. S8**). Patients with core binding factor rearrangements tend to have more mutations than expected for their age (p<10^−15^), yet more favorable outcomes. The mutational spectrum of coding SNVs (**Fig. S5**) accumulates C→T transitions with age (p<10^−3^), with additional C→A transversions in t(8;21) (p<10^−2^) and aberrant karyotype (p<10^−2^) patients.

After adjustment for cytogenetics and multiple comparisons, *NRAS* (p<10^−3^) and *WT1* (p<10^−3^) are mutated significantly more often in younger patients, while *DNMT3A* (p<10^−23^), *IDH1/2* (p<10^−4^), *RUNX1* (p<10^−4^), *TP53* (p<10^−4^), and *NPM1* (p<0.03) are mutated significantly more frequently in older patients. *KRAS, CBL, GATA2, SETD2,* and *PTPN11* mutations appear to be more common in younger patients (0.05<p<0.1, adjusted, **Figs. S7** and **S9**). We identified a prominent hotspot of *MYC* alterations^11^ and previously unreported internal tandem duplications appearing exclusively in children (**Fig. S7**). These observations are replicated in an independent ECOG cohort (**Fig. S10a**) of 384 adult AML patients^5^. Since gene fusions have characteristic cooperating mutations^12^, we devised a weighted resampling scheme to compare mutation frequencies in 584 TARGET and 131 TCGA AML cases while controlling for karyotypic associations. The results (**Fig. S10b**) confirm the generality of the pediatric-adult differences identified above.

For genes such as *CBL, GATA2, WT1, MYC* and *FLT3,* both the frequency and the sites of mutation often differ between children and adults (Figs. 3a and **S7**), with multiple–frequently recurrent-alterations distinct from those identified in adult AML. RAS-related mutations (mutant *KRAS, NRAS, PTPN11,* or *NF1*) are common, particularly with *KMT2A* fusions (**Fig. S11**, **Tables S4-S6**). In addition to being more common and varied, *WT1* mutations appear more likely to be of clonal origin in younger patients (Fig. 3b) despite the majority of pediatric patients presenting with multiple detectable sub-clones (**Fig. S12**).

**Figure 3.**
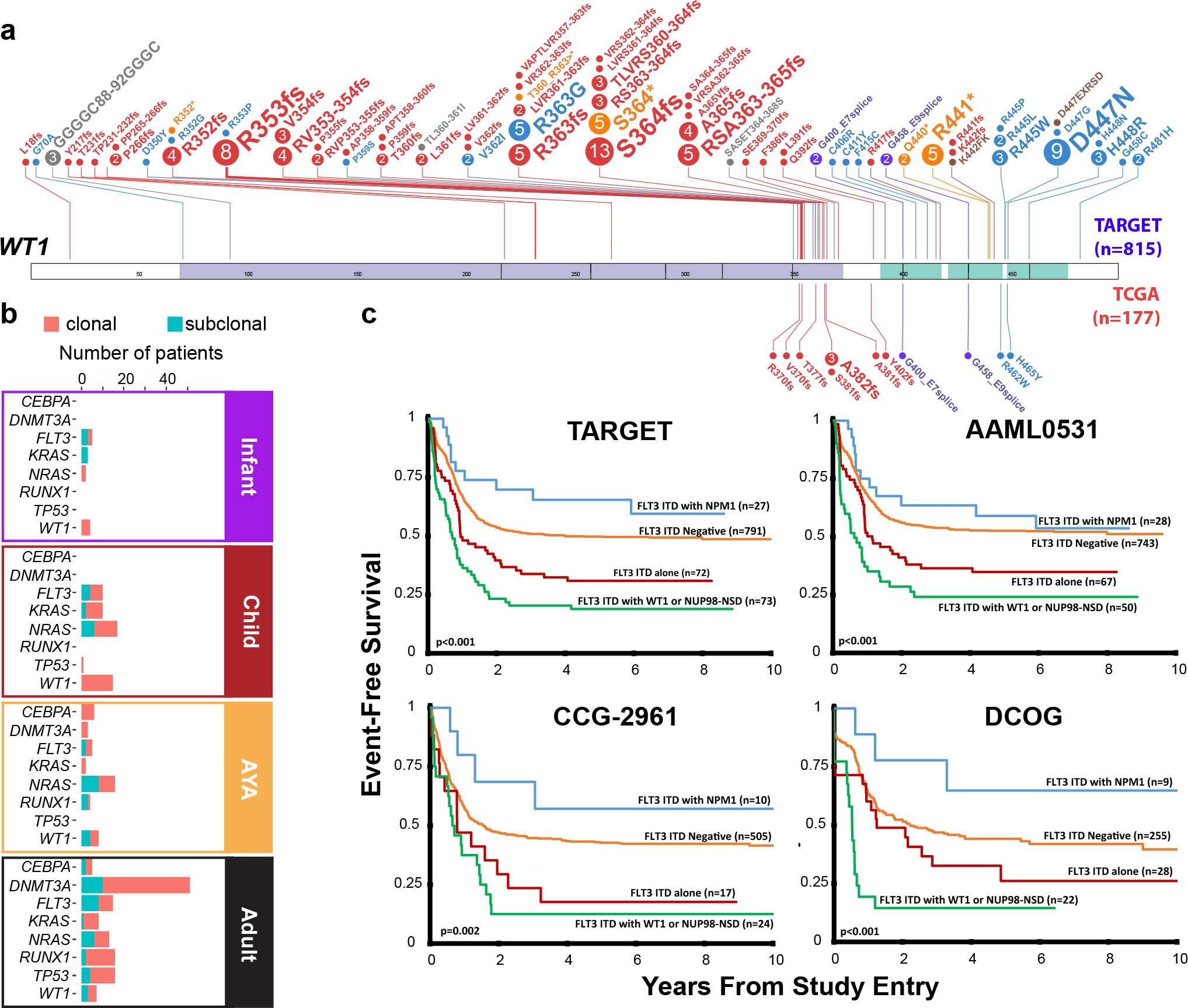
Biological and prognostic interactions between alterations of *WT1, NPM1, FLT3-ITD* and *NUP98-NSD1*. (**a**) *WT1* mutations appear more frequently and impact novel sites in childhood AML (TARGET, expanded above the representation of *WT1* : 18.4%, 150 alterations among 815 patients; TCGA, expanded beneath *WT1* : 7.3%, 13 alterations among 177 patients; Fisher’s exact p = 0.0002). Circles indicate sites of mutation with size proportional to the number of recurrently detected alterations (Colors indicate type of mutation: red, frameshifting; blue, missense; yellow, nonsense; purple, splice site; grey, in-frame deletions; and brown; in-frame insertions. (**b**) Inference of the clonal origin of selected mutations in 197 TARGET AML (Infant, Child and AYA) cases with WGS and 177 TCGA AML (Adult) cases. See Clonality Estimation section in the Online Methods for more details on how the analysis was performed‥ (**c**) The clinical impact of *FLT3*-ITD is modulated by other sequence aberrations. 963 TARGET patients had complete data for *FLT3*-ITD, *NPM1* and *WT1* mutation and *NUP98-NSD1* fusions. Patients with *FLT3*-ITD plus *WT1* and/or *NUP98-NSD1* fusion (n=73) exhibit markedly inferior event-free (multivariate p<0.001) and overall survival (see **Fig. S13**), while co-occurrence of *NPM1* mutations with *FLT3*-ITD associates with improved survival. These findings are confirmed by two separate studies from which TARGET cases were selected (AAML0531 and CCG-2961) as well as an independent cohort of patients treated on European cooperative group trials (DCOG, see online methods).

These differences are clinically significant: we have previously shown that novel *FLT3* mutations are functional, and yield poor responses to standard therapy^13^. The established adverse impact of *FLT3-ITDs* on survival is significantly modulated by co-occurring variants, including *WT1* and *NPM1* mutations and *NUP98* translocations. As shown in Figs. 3c and **S13-S14**, three independent, large-scale studies demonstrate that *FLT3*-ITD accompanied by *NPM1* mutations is associated with relatively favorable outcomes in pediatric patients, while *FLT3*-ITD with *WT1* mutations and/or *NUP98-NSD1* fusions yields poorer outcomes than *FLT3*-ITD alone.

We found no coding mutations in *DNMT3A* in pediatric AML, despite its high frequency in adults. Spontaneous deamination of 5-methylcytosine is strongly associated with aging, and *DNMT3A* contains a CpG dinucleotide yielding hotspot R882 mutations by C-to-T deamination^14^. *DNMT3A* also directly interacts with TP53^15^, itself impacted far more frequently in adults. Mutations of *DNMT3A* or *TP53* drive clonal hematopoiesis in many apparently healthy adults^16^ but are rare in children, as are the *IDH1* and *IDH2* mutations with which they often co-occur.

### The spectrum of somatic structural DNA changes in pediatric AML

Many pediatric AML cases harbor chromosomal copy number changes distinct from those reported in adults (Fig. 4a). Among the 197 cases assayed by WGS, we identified 14 novel focal deletions involving *MBNL1*, a splicing regulator, or *ZEB2*, a key regulator of normal^17^ and leukemic^18^ hematopoiesis (**Fig. S15**). Despite occurring on separate chromosomes, in regions devoid of other deletions, *MBNL1:ZEB2* co-deletions occur far more often than expected (p<10^−13^). Half of these accompany *KMT2A-MLLT3* fusions (p=0.035, **Fig. S11**, **Tables S5-S6**). Samples with *MBNL1:ZEB2* co-deletions carry a larger number of recurrent mutations (p=0.015), and *KMT2A*-fusion samples with del(MBNL1) or del(*ZEB2*) have a larger number of additional cytogenetic abnormalities (p<0.0005). Another 15 novel, validated focal deletions specifically impact *ELF1*, an ETS-family transcriptional regulator of MEIS1^19^. A statistically significant difference in *ELF1* mRNA expression exists between *ELF1*-deleted and intact samples (p<0.01), with 63 genes differentially expressed between the two groups (p<0.01, **Fig. S16**). Among other novel recurrent copy losses, we note five heterozygous deletions of a region containing the *IL9R* gene (**Table S5**) co-occurring with *KIT* mutations and t(8;21).

**Figure 4.**
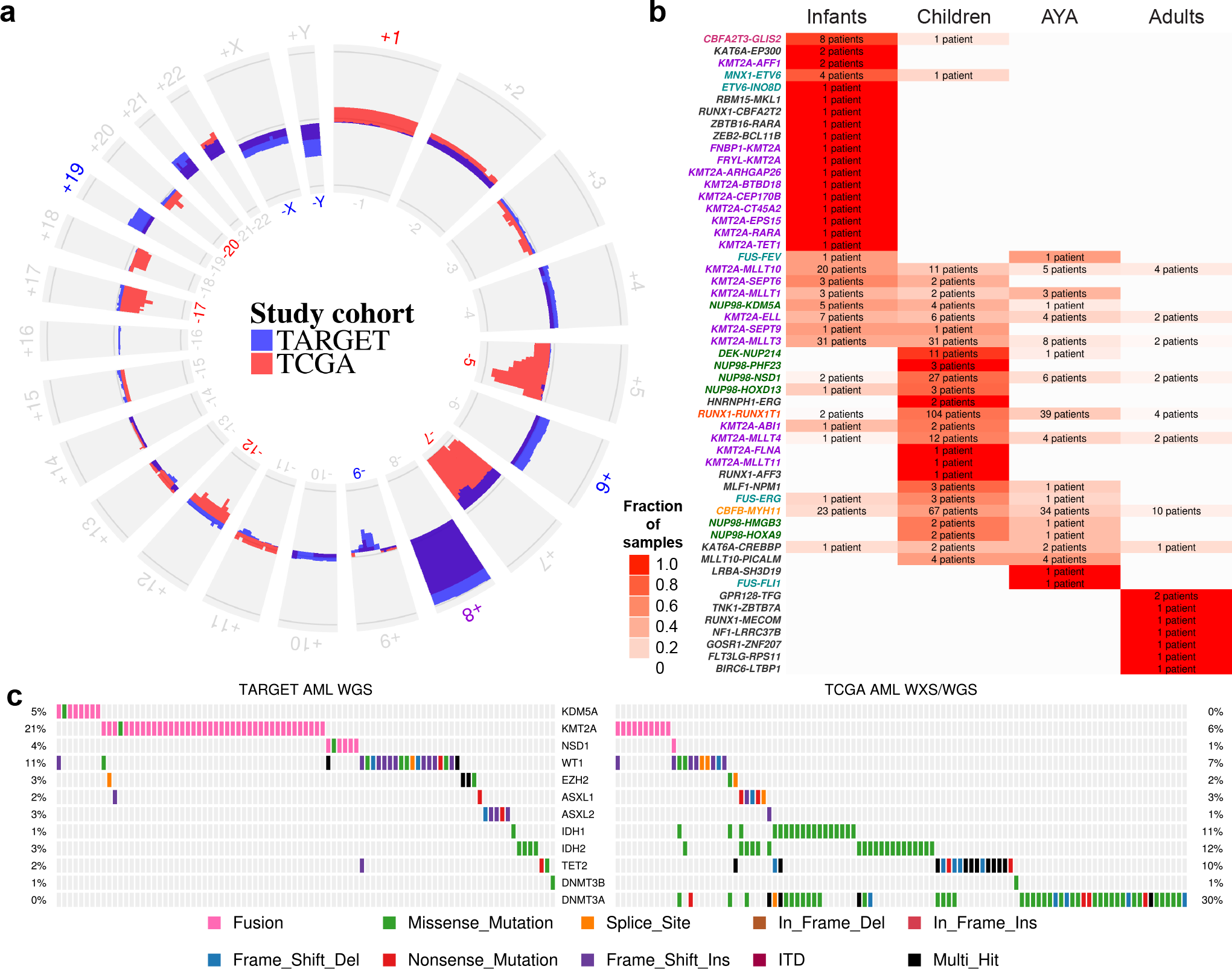
Chromosomal alterations in pediatric and adult AML patients. (**a**) Patterns of regional and chromosomal gain (outward projection) and loss (inward projection) in the TARGET (blue) and TCGA (red) AML cohorts. Losses of 5q, 7, and 17 predominate in adults, while gains of 4, 6, 19, and losses of 9, X, and Y are more common in younger patients. Chromosome numbers are printed on the outside and inside of the circle plot, and colored where there are large pediatric-adult differences. (**b**) Age-specific distributions of validated gene fusions. The fraction of events within an age group for each fusion pair is indicated by white-red shading, while the color of the fusion labels indicates the primary cytogenetic group (colors same as in Fig. 1a, see also **Figs. S17-S18**). The number in each box indicates the number of patients carrying the indicated translocation (labels at left). (**c**) Structural and mutational aberrations affecting epigenetic regulators in TARGET (WGS) and TCGA AML cohorts.

Consistent with our previous findings regarding *NUP98-NSD1* fusions^20^, an expansive catalog of gene fusions, many observed primarily or exclusively in pediatric cases, underscores the disproportionate impact of structural variants in younger patients (Figs. 4b and **S17-S18**). But patterns of exclusion and cooperation are not limited to patients with recurrent structural alteration: mutant *GATA2* is frequently seen in children with normal karyotype (NK) AML, and both *GATA2* (p<10^−9^) and *CSF3R* (p<10^−6^, **Fig. S19**) mutations co-occur with mutations of *CEBPA*^21^. *GATA2* and *CEBPA* are key regulators of hematopoiesis^22,23^, both interacting with *RUNX1* in normal hematopoiesis and leukemogenesis^24^. As with *FLT3/NUP98-NSD1/WT1* interactions, these findings show prognostic interactions in pediatric AML outcome (**Fig. S19b**). *RUNX1* mutations and *RUNX1-RUNX1T1* gene fusions are significantly exclusive of *GATA2* and *CEBPA* mutations (p=0.006, **Fig. S20**, **Table S7**). All four are significantly exclusive of *KMT2A* rearrangements (p<10^−15^), *CBFB-MYH11* gene fusions (p<10^−11^), and *ETV6* aberrations (p=0.01).

### DNA methylation subtypes in pediatric AML

As summarized in Fig. 4c, aberrations affecting epigenetic regulators are widespread and rarely overlap in AML, but their origin (structural vs. mutational) and frequency differs between children and adults. Combining DNA methylation and mRNA expression results in 456 TARGET and TCGA AML cases, we identified dozens of genes with recurrent transcriptional silencing via promoter hypermethylation across TARGET and TCGA AML patients (Figs. 5a **and** 5c, **Tables S8-S9**, details in **Figs. S21-S22**). A number of samples exhibited widespread silencing of genes by aberrant promoter hypermethylation, and this group is enriched for younger patients with *WT1* mutations (p=0.0012, Fig. 5a, hyper-silenced group). Aberrant Wnt/β-catenin signaling is required for the development of leukemic stem cells^25^, and one or more of the Wnt pathway regulators *DKK1, SMAD1, SMAD5, SFRP4, SFRP5, AXIN2, WIF1, FZD3, HES1,* or *TLE1* is deleted or aberrantly methylated in most AML cases^26^. Repression of activating NK cell ligands (particularly *ULBP1/2/3*) appears to be common in pediatric patients, which may represent a therapeutic target^27^. In *KMT2A*-rearranged patients, a cluster of poorly characterized zinc finger genes on chromosome 19 is recurrently silenced.

**Figure 5.**
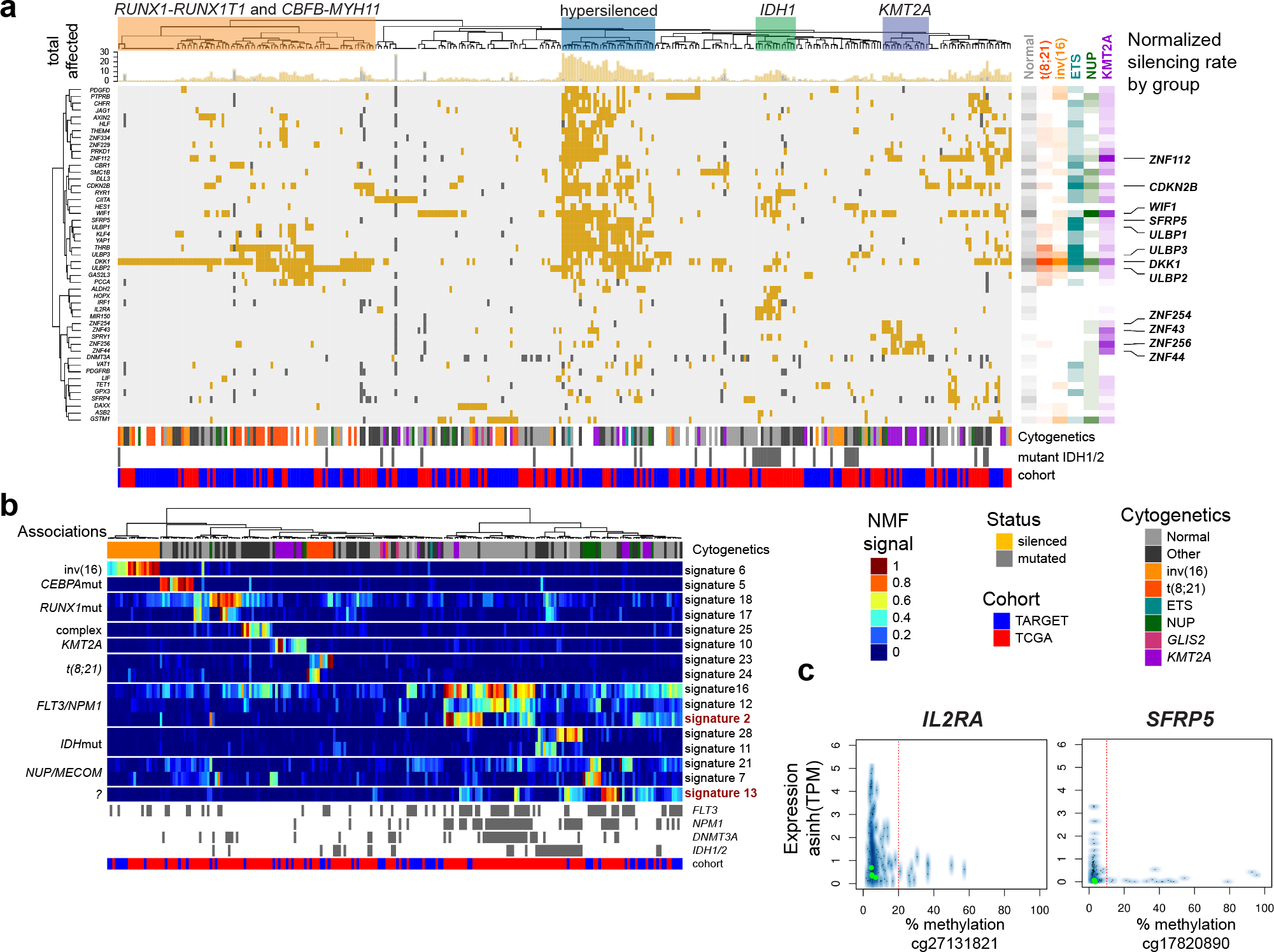
Aberrant DNA methylation in adult and pediatric AML. (**a**) Integrative analysis of genes with recurrent mutations, deletions, or transcriptional silencing by promoter DNA hypermethylation (rows) in TARGET and TCGA AML cases (columns). Cluster associations are labeled at the top, including a prominent group enriched for younger patients with *WT1* mutations (p=0.0012) that shows extensive transcriptional silencing across dozens of genes (blue boxed region, Hypersilenced). The cytogenetic group, *IDH1/2* mutation status (gray, mutated; white, wild-type or unknown) and TARGET/TCGA cohort membership for each sample is indicated below the main figure. The top marginal histogram indicates the total number of genes impacted for each patient. Gene/cytogenetic associations are shown to the right of the main figure, where per gene-rate of involvement by cytogenetic class is indicated by color and shading (unfilled = no involvement; full shading = maximum observed involvement of any gene within patients of the indicated cytogenetic grouping). Wnt regulators and activating NK cell ligands (e.g. *DKK1, WIF1* and *ULBP1, ULBP2, ULBP3,* respectively) are silenced across cytogenetic subtypes (labeled at far right). Distinct groups of silenced genes are also associated with *IDH1* or *IDH2* mutant patients and in KMT2A-rearranged patients. A subset of genes (56 of 119) altered in >3 patients and of patients (n=310; 168 TARGET, 142 TCGA subjects) with one or more genes silenced by promoter methylation is illustrated (see **Figs. S21-S22** and **Tables S8-S9** for enumeration of all 119 genes in all 456 evaluable subjects.). (**b**) A subset (16 of 31) of DNA methylation signatures derived by non-negative matrix factorization (NMF) and *in silico* purification, with samples ordered by hierarchical clustering of signatures (labeled at right). Genomic associations are indicated to the left of the main panel. Signature 13 does not correspond directly to known recurrent alterations, however, along with signature 2 displays potential prognostic significance (see **Fig. S24**). The patient-specific score matrix and display of all 31 signatures are provided in **Table S10** and **Fig. S23**. (**c**) Examples of expression/promoter DNA methylation relationships for *IL2RA* and *SFRP5,* 2 genes identified as recurrently silenced (panel **a**) which also contribute to NMF signatures (panel **b**) are shown. Y-axis: transformed expression (asinh(TPM)), x-axis: promoter CpG methylation. The vertical red line indicates the empirically established silencing threshold.

We applied non-negative matrix factorization (NMF) to CpG methylation data from 284 TARGET and TCGA AML patients with DNA methylation data. By cross-validation, we identified 31 signatures (**Table S10**) that best captured DNA methylation differences across samples, after controlling via *in silico* purification for differences in cellularity. Unsupervised clustering of the resulting DNA methylation signatures largely separated patients by age and karyotypic subtypes (Figs. 5b and **S23**), but also revealed a signature which did not associate strongly with age or established prognostic factors (Signature 13, Fig. 5b). Two signatures (signatures 2 and 13) predicted significantly (p < 0.05) poorer event-free survival in both pediatric and adult patients with above-median scores, after stratifying by cohort and adjusting for *TP53* mutation status and white blood cell count (**Fig. S24**). Larger sample sizes are needed to evaluate the clinical significance of these findings.

### The pediatric AML transcriptome is shaped by diverse miRNAs

We performed miRNA sequencing of 152 cases to characterize miRNA expression patterns in pediatric AML. Unsupervised clustering of the data revealed 4 discrete subgroups that were correlated with specific genomic alterations (Figs. 6a and **S25**), including high miR-10a expression in samples with *NPM1* mutations, consistent with previous reports^28^. Further, Cox proportional hazards analyses identified multiple miRNAs associated with clinical outcome (**Figs. S26-S28**, **Table S11**), including miR-155, which we previously reported to predict poor survival^29^.

**Figure 6.**
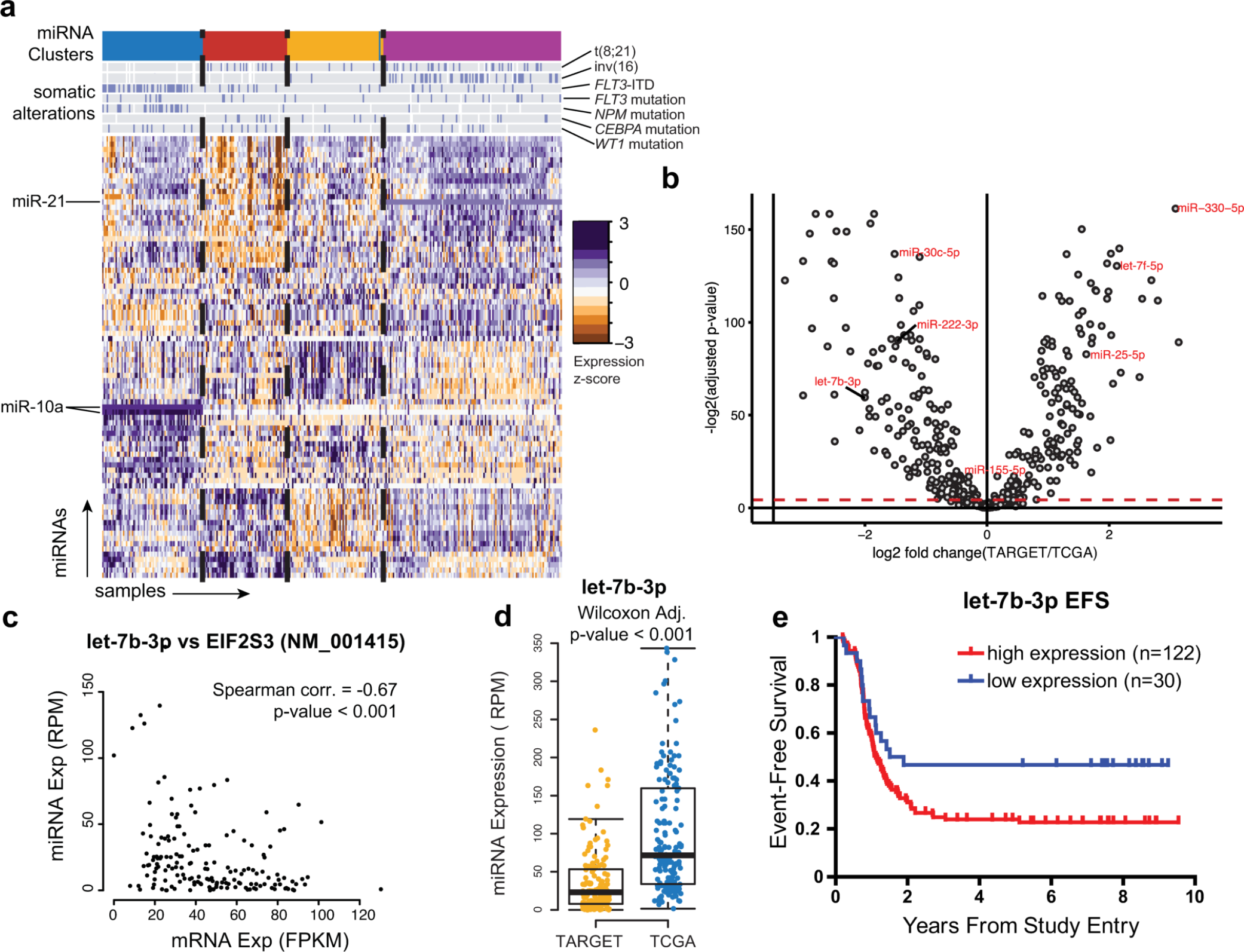
miRNAs differentially regulate distinct molecular and age sub-groups in AML. (**a**) Unsupervised clustering of miRNA expression patterns in 152 childhood AML cases identifies four patient subgroups (colored bands at top) with correlation to somatic alterations as indicated (blue bars on gray background), and subgroup-specific miRNA expression (miR-10 and miR-21 are highlighted as examples). (**b**) Age-related differences in miRNA expression are evident between adult (n=162) and pediatric AML (n=152). Volcano plot indicates differentially expressed miRNAs between adult and pediatric cases. Red-green point shading indicates relative under- or over-expression in TARGET, respectively. (Wilcoxon test, Benjamini-Hochberg adjusted P<0.05; Threshold indicated by dashed red line). (**c**) A predicted miRNA:mRNA target relationship involving *let-7b,* which is (**d**) less abundant in most pediatric cases than in adult cases. (**e**) High expression of *let-7b* occurs in a minority of pediatric AML and is associated with shorter time to relapse.

Differential expression analyses using Wilcoxon tests revealed miRNAs that are differentially expressed between pediatric and adult AML (Fig. 6b). Of note, miR-330 was the most over-expressed in pediatric samples, and has previously been shown to have oncogenic potential in AML^30^.

Several age-associated miRNAs harbor binding sites within, and have expression levels anti-correlated with, putative target genes that may be involved in RNA and protein processing suggesting that miRNAs could contribute to leukemogenesis through the dysregulation of transcripts and proteins^31^. Of note, *let-7b,* which is a potential regulator of protein synthesis via *EIF2S3* (Fig. 6c), is typically less abundantly expressed in pediatric AML (Fig. 6d). However, high *let-7b* expression in pediatric AML is associated with shorter time to relapse (log-rank p<0.05, Fig. 6e).

### Discussion

Using a large cohort of patients, this study establishes the prevalence of, and coincident relationships among, recurrent somatic genetic and epigenetic abnormalities in pediatric AML. We observe several features in common between pediatric and adult AML: a low overall mutation rate in comparison to other cancers, a long tail of infrequently affected genes, and overlap among recurrently impacted genes. However, pediatric AML exhibits distinctive and critically important characteristics. We and others have previously reported on the presence and clinical impact of novel fusion genes in pediatric AML^20,32^. As this study illustrates, the impact of fusion transcripts in AML is both broad and age-dependent. Recognition and comprehensive testing for these alterations are key first steps in the development of new and potentially novel modes of targeted therapy^33^.

Recurrent focal deletions represent a unique aspect of pediatric AML. Regional (e.g. chromosomal arm-and band-level) copy loss differs substantially by age, but surprisingly, focal areas of copy loss are also more common in children, specifically impacting *ZEB2, MBNL1,* and *ELF1*. *MBNL1* is upregulated by the *KMT2A-AF9* fusion protein^34^, and genes involved in post-transcriptional processing (*SETD2, U2AF1, DICER1*) harbor the sole recurrent mutation in several *KMT2A*-rearranged cases, suggesting a functional role for altered splicing in pediatric leukemogenesis. Alterations in *ZEB2* have been identified as cooperating events in murine *CALM-AF10* leukemia models^35^ while *ZEB2* knockout mice develop myelofibrosis^36^, suggesting a fundamental role for this gene in the pathogenesis of AML.

Many of the genes characteristically mutated in AML are altered at widely variable frequencies across age groups; several (including *FLT3* and *WT1*) are impacted by pediatric specific variants and hotspots. Clinical tests for a handful of genomic alterations are widely used to risk-stratify patients and determine treatment regimens. However, the current practice of considering the effect of each somatic alteration in isolation is inadequate. As we illustrate for *FLT3*-ITD, interactions among sequence variants can have dramatic clinical consequences. Moreover, some interactions appear to be age-specific. In pediatric AML, *FLT3*-ITD and *NPM1* mutations co-occur in the absence of *DNMT3A* mutations in a group of patients with superior outcomes (Figs. 3c, **S13** and **S14**), in contrast to inferior outcomes reported in adults where *FLT3*-ITD and *NPM1* mutations frequently co-occur with mutations in *DNMT3A*^4^. In the TCGA adult AML cohort, over half the subjects with somatic *FLT3* and *NPM1* mutations also possessed somatic *DNMT3A* mutations^3^. Subsequent studies established the generality of this result^4^, and revealed that *DNMT3A* mutations are early clonal events^37^, which often co-operate with later *NPM1* and *FLT3* mutations to promote chemoresistance, mutagenesis,^38^ and inferior outcomes^39^. Similarly, the co-occurrence of *FLT3*-ITD with *WT1* mutations or *NUP98-NSD1* fusions accompanies frequent induction failure and dismal outcomes in children with AML (multivariate p<10^−4^, Figs. 3c, **S13** and **S14**).

In TARGET, TCGA, and ECOG AML cases, *WT1* mutations were mutually exclusive with those of *ASXL1* and *EZH2* (p < 10^−3^). *WT1* recruits *EZH2* to specific targets^40^, and *WT1* mutations have been linked to promoter DNA hypermethylation of *EZH2* target genes^41^. Mutant *ASXL1* abolishes *EZH2*-mediated silencing of HOX genes^42^. *EZH2* resides on a recurrently deleted region of chromosome 7, and decreased *EZH2* activity is associated with treatment resistant AML^43^. In pediatric AML, mutant *WT1* and *EZH2* appear to be of exclusively clonal or near-clonal origin, with nearly a quarter of TARGET cases harboring mutations affecting one or the other. Aberrant *WT1*, *EZH2*, or *ASXL1* predicted induction failure in TARGET AML cases (multivariate p<0.05, adjusted for interactions with *FLT3* alterations, *NUP98-NSD1,* and *KMT2A* fusions) and were largely mutually exclusive with *KMT2A* rearrangements (p < 10^−5^). Many of these patients present without apparent chromosomal abnormalities at diagnosis, yet less than 20% achieve long-term remission with standard treatment, highlighting the importance of molecular stratification to achieve better outcomes. It is possible that early events such as *WT1* mutations and *NUP98-NSD1* fusions in children may play a similar role to that observed for *DNMT3A* mutations^14^ in adults, with significant implications for risk stratification in AML across age groups.

Our data also demonstrate that DNA-methylation and miRNA expression profiles both accompany and complement DNA alterations, and can stratify pediatric AML patients in terms of both overall and progression-free survival. These findings suggest a need to update pediatric AML clinical risk categories beyond current classifications, with important implications for clinical practice.

Despite incremental improvements with increasingly intensified regimens, modern outcomes in pediatric AML have plateaued, with only ^~^60% of patients achieving long term survival. As many as 10% of children will die from direct complications of treatment. Survivors suffer unacceptably high rates of long-term morbidities resulting from anthracycline exposure or sequelae of hematopoietic stem cell transplantation. As illustrated herein, pediatric AML is a collection of molecularly diverse diseases with similar phenotypes. No single treatment strategy is likely to be effective for all pediatric AML subtypes, which may explain repeated failures of randomized clinical trials to improve outcomes in recent years. In keeping with the shift towards comprehensive, molecularly based classification schemas in AML^4^, the time has come to develop targeted therapies that address specific vulnerabilities of pediatric subtypes. The TARGET AML dataset will serve as a foundation for development of pediatric-specific classification schemas and the development of personalized treatment strategies.

## Acknowledgements

Dedicated to the memory of our colleague, mentor and friend, Dr. Robert Arceci, whose vision and perseverance set this effort in motion: “I may not have gone where I intended to go, but I think I have ended up where I needed to be.” (Douglas Adams, *The Long Dark Tea-Time of the Soul*)

The results published here are based upon data generated by the Therapeutically Applicable Research to Generate Effective Treatments (TARGET) initiative and The Cancer Genome Atlas. Data used for this analysis are available under dbGaP accession numbers phs000465 and phs000178. The TARGET initiative is supported by NCI Grant U10CA98543. Work performed under contracts from the National Cancer Institute, US National Institutes of Health within HHSN261200800001E includes specimen processing (the Children’s Oncology Group Biopathology Center), whole genomic sequencing (Complete Genomics) and RNA and targeted capture sequencing (British Columbia Cancer Agency). The content of this publication does not necessarily reflect the views or policies of the Department of Health and Human Services, nor does mention of trade names, commercial products, or organizations imply endorsement by the U.S. Government. Computation for the work described in this paper was supported in part by Fred Hutchinson Scientific Computing, the University of Southern California’s Center for High-Performance Computing, and NSF award ACI-1341935. This work was additionally supported by COG Chairs U10CA180886 and U10CA98543; COG Statistics and Data Center U10CA098413 and U10CA180899; COG Specimen Banking U24CA114766; R01CA114563 (SM); St. Baldrick’s Foundation (JEF, TT, SM); Alex’s Lemonade Stand (SM), Target Pediatric AML (TpAML), P20GM121293 (JEF); Arkansas Biosciences Institute (JEF), and the Jane Anne Nohl Hematology Research Fund (TT).

## Author Contributions

HB, JEF, TT and RER contributed equally to this work. MAS, DSG, SM and RA (see Acknowledgements) conceived and led the project. RER, MAM, JMGA, TMD, PG, LCH, DSG and SM managed the project. HB, JEF, TT, RER, ELL, TAA, YM, RM, AJM, MAM, JZ, XM, YuL, YaL, TMD, ACH, BS, and SRP generated, processed, and analyzed the data. SC, GR, CMZ, SN, EAK and ASG shared critical data and reagents. HB, JEF, TT, RER, ELL and SM drafted the manuscript. All authors edited and approved the manuscript.

## Online Methods

### Sample Selection and Preparation

All patient samples were obtained by member COG institutions after written consent from the parents/guardians of minors upon enrolling in the trial. The study was overseen by the Institutional Review Board at Fred Hutchinson Cancer Research Center (Protocol 1642, IR File #5236). Selected clinical (e.g., age, presenting hematological indices, cytogenetic classification) and molecular features (e.g., *KIT, RAS, NPM, WT1, CEBPA, IDH1* mutations, and *FLT3*/ITD allelic ratios) were clinically available prior to genomic analyses and are included in the clinical data file available at the TARGET data matrix. 177 cases from the adult de novo AML TCGA dataset^3^ were selected for analysis after exclusion of those with FAB M3 morphology (n=20) or *BCR-ABL1* gene fusion (n=3) since these subtypes are not represented in the COG/TARGET-AML cohort. The age distributions for the TARGET WGS discovery group and the TCGA cohort are outlined in **Table S3**.

DNA and RNA was extracted from ficoll-enriched, viably cryopreserved samples from the COG biorepository using the AllPrep Extraction Kit (Qiagen). Nucleic acids were quantified by NanoDrop (Thermo Scientific). RNA samples were tested for quality and integrity using the Agilent 2100 Bioanalyzer (Agilent Technologies). The integrity of DNA samples was confirmed by visualization on a 0.8% agarose gel.

### Whole genome sequencing

Sequencing libraries were constructed for WGS cases from genomic DNA and sequenced using combinatorial probe anchor ligation by Complete Genomics (CGI)^44^. Reads were mapped to the GRCh37 reference human genome assembly by the CGI Cancer Sequencing service using software version 2.1 of the CGI cancer analysis pipeline (http://www.completegenomics.com/customer-support/documentation/).

Somatic coding SNVs and indels were extracted from MAF files and filtered to remove 1) germline variants; 2) low-confidence variants and 3) paralogs. For step 1, germline variants used for filtering include those from NLHBI Exome Sequencing Project (http://evs.gs.washington.edu/EVS/), dbSNP 132 (https://www.ncbi.nlm.nih.gov/projects/SNP/), St Jude/Washington University Pediatric Cancer Genome Project (PCGP), and CGI WGS from the TARGET project. For step 2, a mutation is considered of low-confidence if it does NOT meet one of the following criteria: a) mutant allele has ≥3 more read count in tumor than in the normal sample; b) the mutant read count in tumor is significantly higher than that in the matched normal (P<0.01 by Fisher exact test); and c) mutant allele fraction in normal is below 0.05. For step 3, we ran BLAT search using a template sequence that includes the mutant allele and its 20-bp flanking region to determining the uniqueness of mapping of the mutation. To avoid over-filtering, we implemented a rescue pipeline which retains all “gold” variants that match known somatic mutation hotspots based on our variant classification program Medal Ceremony^45^.

In addition to small variant calls (SNV, indel), the CGI cancer analysis pipeline delivered flat files of potential novel DNA junctions and segmented copy number ratios derived from normalized read counts from paired tumor/normal specimens. Circos summary plots of the unfiltered CGI data are available through the data matrix. To reduce potentially spurious calls, final CNVs for analysis were trimmed after empirical tuning to previously available Affymetrix SNP6 microarray calls in matched samples by requiring a CGI average normalized coverage (avgNormCvg) in the region of ≥20 for putative non-homozygous deletions, the SD for lesser allele fraction ≤ 0.22, a CGI ploidyScore of <30 and trimming of calls on ChrM, centromeric or telomeric regions, and merged for adjacent CNV, per called direction, within 10 Kbp. With these filters, 75 and 85% (loss and gain, respectively) of filtered CNV calls matched CNVs previously called by Affymetrix SNP6 microarray and 87% of chromosome-arm level calls matched reported karyotype abnormalities reported in the clinical data. Putative copy variants underwent further secondary confirmation using the nanoString nCounter assay (Nanostring Technologies). Novel DNA junctions discovered by WGS were included in cases where at least one additional level of support was available, either from cytogenetic analysis or from RNA sequencing studies.

### Targeted Capture Sequencing

Candidate genes identified by WGS analysis were selected for independent verification in 182 samples from the WGS discovery cohort and 618 additional subjects treated on COG AAML0531. Capture baits were designed and ordered using Agilent’s SureDesign (https://earray.chem.agilent.com/suredesign/) for these selected genes along with target regions identified in concurrent TARGET studies, targeting coding regions and UTRs with a 10 bp pad. This design (TARGET AML + TARGET other) resulted in an overall target space of 2.376 Mbp with 98.7% of target regions covered by a probe. Probe density was specified at 2x, with moderately stringent repeat masking, and balanced boosting options selected.

Genomic DNA libraries from which gene regions of interest are captured were constructed according to British Columbia Cancer Agency Genome Sciences Centre (BCGSC) plate-based and paired-end library protocols on a Biomek FX liquid handling robot (Beckman-Coulter, USA). Briefly, 1μg of high molecular weight genomic DNA was sonicated (Covaris E210) in a 60μL volume to 200-300bp. Sonicated DNA was purified with magnetic beads (Agencourt, Ampure). The DNA fragments were end-repaired, phosphorylated and bead purified in preparation for A-tailing. Illumina sequencing adapters were ligated overnight at 20°C and adapter ligated products bead purified and enriched with 4 cycles of PCR using primers containing a hexamer index that enables library pooling. 94ng from each of 19 to 24 different libraries were pooled prior to custom capture using Agilent SureSelect XT Custom 0.5-2.9Mb probes. The pooled libraries were hybridized to the RNA probes at 65°C for 24 hours. Following hybridization, streptavidin-coated magnetic beads (Dynal, MyOne) were used for custom capture. Postcapture material was purified on MinElute columns (Qiagen) followed by post-capture enrichment with 10 cycles of PCR using primers that maintain the library-specific indices. Paired-end 100 base reads were sequenced per pool in a single lane of an Illumina HiSeq2500 instrument. Illumina paired-end sequencing reads were aligned to the GRCh37-lite reference using BWA version 0.5.7. This reference contains chromosomes 1-22, X, Y, MT, 20 unlocalized scaffolds and 39 unplaced scaffolds. Multiple lanes of sequences were merged and duplicated reads were marked with Picard Tools. Small variants (SNV and indel) from TCS data were identified by parallel methods, integrated, and subsequently filtered as follows. **Mpileup**: SNVs were analyzed with SAMtools mpileup v.0.1.17 on paired libraries^46^. Each chromosome was analyzed separately using the -C50-DSBuf parameters. The resulting vcf files were merged and filtered to remove low quality variants by using samtools varFilter (with default parameters) as well as to remove variants with a QUAL score of less than 20 (vcf column 6). Finally, variants were annotated with gene annotations from ensembl v66 using snpEff^47^ and the dbSNP v137 db membership assigned using snpSift^48^. **Strelka**: Samples were analyzed pair wise with the default settings of Strelka^49^ v0.4.7 with primary tumor samples against the matched remission sample. Somatic variants called by either Mpileup or Strelka were combined and filtered by meeting any of the following criteria: <10 reads in the remission sample, <10 reads in the tumor sample, tumor alt base = 0, adjusted tumor allele frequency = 0, gmaf >0.009, or >60 patients had exact SNV. For patients established to be in morphological remission, additional filters included removing variants with >0.10 allele fraction in the remission sample and a FET score of >0.05. For refractory patients, variants were excluded with >0.35 allele fraction in the post-Diagnostic sample. These filtered variants could be “rescued” if a variant was a known COSMIC mutation associated with hematological cancers. The filtering criteria for indel calls were similar. Tandem duplications were identified with Pindel using default parameters^50^. In addition, clinical molecular testing for specific genes (*FLT3* ITD and *FLT3* codons 835/836, *CEBPA* bzip and NTD regions, *KIT* exons 8 and 17, *CBL* exons 8 and 9, and *WT1* exon 7) were merged into the variant calls for final analysis.

DNA variants from discovery and TCS studies were merged to construct the mutation profile for each gene using the web-based program, ProteinPaint^51^. Genome-wide mutational burden was compared to published data from, and using the method of, Lawrence et al^52^.

### *CBL* Transcript Variant Screening by cDNA PCR

Total RNA isolated from patient leukemic cells using the Qiagen AllPrep DNA/RNA Mini Kit (Qiagen, Germany) was reverse transcribed to cDNA with oligo DT primer and additional reagents following the Maxima H Minus First Strand cDNA Synthesis Kit instructions (Thermo Scientific, Grand Island, NY).

Synthesis of the second-strand cDNA and following PCR were performed using the following primers: forward primer for genemap: (5′FAM-TTCCAAGCACTGATTGATGG), forward primer for sequencing (5′-TTCCAAGCACTGATTGATGG-3′), reverse primer: (5′-AACAGAATATGGCCGGTCTG). PCR was performed in 25uL volumes containing 12.5uL Failsafe Epicentre Buffer C (2x) (Epicentre Technologies, Madison, WI), 0.5uL (10uM) of each primer, 0.25uL Invitrogen Platinum Taq Polymerase (Thermo Scientific - Invitrogen, Grand Island, NY), 1uL of cDNA, and 10.25uL of Nuclease-Free Water (USB Corporation, Cleveland, OH). The Thermocycling program consisted of 5 min denaturation at 95 C, followed by 35 cycles at 95 C for 30 sec, 60°C for 30 sec and 72 C for 45 sec min and a final extension of 7 min at 72 C in a 96 well Biometra T professional Thermocycler (Biometra, Germany)

PCR products were diluted in nuclease free water (USB Corporation, Cleveland, OH) and mixed with deionized Formamide and GENESCAN-400HD (ROX) size markers (Applied Biosystems, Foster City, CA) and submitted for electrophoresis on an ABI 3730 Genetic Analyzer (Applied Biosystems). After electrophoresis the fluorescence signals were analyzed using GeneMapper 5.0 software (Applied Biosystems). Genemapper screening revealed products of the expected WT normal size (685bp), and additional products of various sizes: corresponding to complete deletion of Exon 8 (563bp), complete deletion of Exon 9 (485bp), as well as deletion involving both CBL exon 8 and exon 9 (354bp).

Patients exhibiting deletions by Genemapper were then sent for sequence verification. PCR products were treated with Exo-SAPit enzyme (USB Corporation, Cleveland, OH). Sequencing was done by Eurofins MWG Operon LLC (Huntsville, AL) in accordance with their DNA sequencing process guidelines and methods.

### Generalized linear mixed model for coding mutation counts

In order to account for both fixed and random effects which might be present with age and cytogenetic subgroups, we employed a generalized linear mixed model (glmm, Knudson 2016, R package version 1.1.1, https://CRAN.R-project.org/package=glmm) to model the discrete counts of coding SNVs in each TARGET and TCGA WGS patient with a Poisson error distribution (log link). Marginal likelihood ratio tests for age (as a continuous predictor) and cytogenetic subgroup (as a categorical predictor) were uniformly and highly significant, as reported in the text, while the per-cytogenetic-group random effects accounted for a small (< 0.003%) fraction of the variance observed. The model converged in 208 steps; 10,000 MCMC iterations were employed to estimate the mixed effects component of the model, fitted per-cytogenetic-group assuming a random slopes model.

### Generalized Dirichlet-multinomial regression for mutational spectra

To accommodate the possibility of either negative or positive correlation between the counts of each type of mutation (C→T, C→A, C→G, T→C, T→A, T→G) in each subject, we employed a generalized Dirichlet-multinomial model (mglm^53^, R package version 0.0.7, https://CRAN.R-project.org/package=MGLM) with age and cytogenetic group as predictors, mutational spectrum (a matrix of counts for each type of mutation) as response. At convergence, the significant predictors of mutational spectrum differences were age (most significant), t(8;21) status, and aberrant karyotype (mutually exclusive with t(8;21) and other common recurrent chromosomal abnormalities). C→T transitions are known to increase with age, particularly for methylated cytosines; however, an inflation of C→A transversions was particularly apparent in t(8;21) and aberrant karyotype cases. (Both t(8;21) and inv(16) affect core binding factor subunits, and both are associated with higher mutational burdens at a given age, but only t(8;21) cases show additional C→A transversions beyond those expected from counts).

### Weighted resampling scheme to compare TARGET and TCGA mutation frequencies

Common chromosomal aberrations often co-occur with specific types of additional DNA sequence abnormalities. To account for this observation when determining differences in mutation frequencies between TARGET AML and TCGA AML, we first divided each cohort into the following categories: *KMT2A* fusions, t(8;21), inv(16), del(7), +8, +21, -Y, and normal karyotype (NK). A total of 131 unique TCGA and 548 TARGET samples fell into one of the above categories. We then sampled equal numbers of specimens from each category and calculated the fraction of samples with mutations in a given gene. To account for sampling variations, we repeated our sampling procedure 5000 times and calculated the mean and standard deviation of the fraction of samples with mutations in each gene of interest.

### Variant pairwise mutual exclusion and co-occurrence

Pairwise mutually exclusive sequence alterations (**Fig S20a**) were identified using CoMEt^54^ with the “exhaustive” option (http://compbio.cs.brown.edu/projects/comet/). Pairwise co-occurrence p-values (**Fig S20b**) were calculated directly using a hypergeometric distribution (equivalent to Fisher’s exact test). Statistically significant exclusion/co-occurrence patterns were visualized using Cytoscape^55^ (http://cytoscape.org/), with edge thickness representing–log10(p-value).

### Orthogonal evaluation of mutual exclusion and co-occurrence via penalized Ising model

A slightly different approach to reconstructing a binary-valued undirected graph (a discrete Markov random field) employs penalized logistic regression of all candidate nodes upon each possible target and selects the most probable graph structure based on extensions of the Bayes information criterion (EBIC). This approach is implemented by Epskamp (https://cran.r-project.org/package=IsingFit)^56^ and employs a hyperparameter (γ) for the penalty weight which eventually determines the density of the estimated network. Adjustment for multiple comparisons was applied to the marginal significance of each gene-gene Fisher exact test; this value is not unbiased due to post-selection inference and is only intended as a guide. The resulting network of correlated and anticorrelated binary indicators (gene-and chromosomal-level aberrant/wildtype, pediatric/adult) recovers known and CoMEt-detected relationships, but also identifies several novel and marginally significant (by Fisher’s exact test, see above) relationships, as summarized in Supplementary Table 6.

### Hypothesis-testing

Except where described by the methods above, p-values are calculated by Fisher’s exact test; where an exact binomial test is impractical, we approximate this with a Chi-squared p-value.

### Regression fits for structural/sequence variant burden and age-associated recurrent abnormalities

To fit the ratio of structural to sequence variant impact in each patient, we added 0.333 as a smoothing factor to the counts of each clonal event of each type, using all recurrently mutated, fused, or silenced genes, identified in either cohort, as candidates for “impact” by structural variants. The transparency of each data point represents its observed over expected mutational burden, given the patient’s age, but has no impact on the loess regression fit. The loess curve was fit by ggplot2 (http://ggplot2.org) on a log10 scale. To estimate the relative contribution of each of the recurrent fusion neighborhoods across ages (rather than age groups), we used the “zoo” time series package^57^ to fit a rolling median with expanding time steps (1, 3, 5, 8, 17) across all subjects for whom we had data on fusions. The (smoothed) contribution of each family of fusions to the total number of patients in a given age window (expanding with advancing age) is plotted in Figure 2d.

### Clonality estimation

Several packages (including MAFtools^58^ (https://github.com/PoisonAlien/maftools) Gaussian mixture, SciClone’s^59^ beta mixture model, and a weighted penalized logistic mixture model) were compared to validate the results obtained, in addition to manual review of all results. While proportions of mutations assigned to various clones differed in some cases (especially with and without read support weighting), the primary mutational clones were consistently identified by all methods, and an overall tendency for childhood and AYA patients to present with greater diagnostic mutational clonality, at the read depths available in the TARGET WGS and TCGA data, was confirmed by all methods. Among AYA patients (where both TCGA and TARGET AML cohorts contain numerous patients), no difference in estimated clonality or monoclonal/polyclonal balance was observed between cohorts (p=0.7 and p=0.65 respectively by Fisher’s exact test), and although a trend towards decreased mutational clonality with increasing age among AYA patients was observed, it was not statistically significant (p=0.2). It is important to note that, owing to variable sequencing depths, we do not have the statistical power to reliably detect clones present in less than 5% of the total sample material, though inclusion of variant allele frequencies as low as 0.1% did not change our results or conclusions regarding mutational clonality. Karyotypic clonality was assessed by parsing ISCN karyotypes of all TARGET and TCGA AML patients and using stemline karyotype to identify the most likely ancestral aberrations for patients with abnormal karyotype. Patients with normal karyotype were assigned a karyotypic clonality of 1, as were patients with all metaphases bearing identical aberrations.

### Aberrations predicting induction failure

A logistic model with terms for *NUP98-NSD1* fusions, *FLT3* mutations, interactions between the preceding, and (any one of) *WT1, EZH2,* or *ASXL1* mutation (mutually exclusive) or deletions of the latter (nearly mutually exclusive), or *KMT2A* rearrangements (also mutually exclusive with the preceding) best fit the data for subjects where the first recorded event was either induction failure (1) or any other outcome (0). All possible nested models with the same terms, and all other models arrived at by penalized logistic regression (using an elastic net penalty with the *glmnet* package^60^, with any observed recurrent lesion eligible for inclusion as an independent predictor), yielded inferior fits both in terms of classification error and by Akaike information criterion (AIC). We report the marginal p-value for WT1/ASXL1/EZH2 aberrations as predictors of induction failure in the test based on this model fit.

### mRNA Sequencing

Total RNA quality was verified on Agilent Bioanalyzer RNA nanochip or Caliper GX HT RNA LabChip, with samples passing quality control arrayed into a 96-well plate. PolyA+ RNA was purified using the 96-well MultiMACS mRNA isolation kit on the MultiMACS 96 separator (Miltenyi Biotec) from 2μg total RNA with on-column DNaseI-treatment as per the manufacturer’s instructions. The eluted PolyA+ RNA was ethanol precipitated and resuspended in 10μL of DEPC treated water with 1:20 SuperaseIN (Life Technologies). First-stranded cDNA was synthesized from the purified polyA+RNA using the Superscript cDNA Synthesis kit (Life Technologies) and random hexamer primers at a concentration of 5μM along with a final concentration of 1ug/uL Actinomycin D, followed by Ampure XP SPRI beads on a Biomek FX robot (Beckman-Coulter). The second strand cDNA was synthesized following the Superscript cDNA Synthesis protocol by replacing the dTTP with dUTP in dNTP mix, allowing second strand to be digested using UNG (Uracil-N-Glycosylase, Life Technologies, USA) in the post-adapter ligation reaction and thus achieving strand specificity. The cDNA was quantified by PicoGreen (Life Technologies) and VICTOR^3^V Fluorimeter (PerkinElmer). The cDNA was fragmented by Covaris E210 sonication for 55 seconds at a “Duty cycle” of 20% and “Intensity” of 5. The paired-end sequencing library was prepared following the BC Cancer Agency Genome Sciences Centre strand-specific, plate-based and paired-end library construction protocol on a Biomek FX robot (Beckman-Coulter, USA). Briefly, the cDNA was purified in 96-well format using Ampure XP SPRI beads, and was subject to end-repair, and phosphorylation by T4 DNA polymerase, Klenow DNA Polymerase, and T4 polynucleotide kinase respectively in a single reaction, followed by cleanup using Ampure XP SPRI beads and 3′ A-tailing by Klenow fragment (3′ to 5′ exo minus). After purification using Ampure XP SPRI beads, picogreen quantification was performed to determine the amount of Illumina PE adapters to be used in the next step of adapter ligation reaction. The adapter-ligated products were purified using Ampure XP SPRI beads, and digested with UNG (1U/μl) at 37oC for 30 min followed by deactivation at 95oC for 15 min. The digested cDNA was purified using Ampure XP SPRI beads, and then PCR-amplified with Phusion DNA Polymerase (Thermo Fisher) using Illumina’s PE primer set, with cycle condition 98°C 30sec followed by 10-13 cycles of 98°C 10 sec, 65°C 30 sec and 72°C 30 sec, and then 72°C 5min. The PCR products were purified using Ampure XP SPRI beads, and checked with Caliper LabChip GX for DNA samples using the High Sensitivity Assay (PerkinElmer, Inc. USA). PCR product of the desired size range was purified using 8% PAGE, and the DNA quality was assessed and quantified using an Agilent DNA 1000 series II assay and Quant-iT dsDNA HS Assay Kit using Qubit fluorometer (Invitrogen), then diluted to 8nM. The final library concentration was double checked and determined by Quant-iT dsDNA HS Assay again for Illumina Sequencing.

### mRNA Quantification

Illumina paired-end RNA sequencing reads were aligned to GRCh37-lite genome-plus-junctions reference using BWA version 0.5.7. This reference combined genomic sequences in the GRCh37-lite assembly and exon-exon junction sequences whose corresponding coordinates were defined based on annotations of any transcripts in Ensembl (v69), Refseq and known genes from the UCSC genome browser, which was downloaded on August 19 2010, August 8 2010, and August 19 2010, respectively. Reads that mapped to junction regions were then repositioned back to the genome, and were marked with ’ZJ:Z’ tags. BWA is run using default parameters, except that the option (-s) is included to disable Smith-Waterman alignment. Finally, reads failing the Illumina chastity filter are flagged with a custom script, and duplicated reads were flagged with Picard Tools. Gene, isoform, and exon-level quantification was performed as previously described^61^.

### Fusion mRNA Transcript Detection

Transcriptomic data were de novo assembled using ABySS (v1.3.2) and trans-ABySS (v1.4.6)^62^. For RNA-seq assembly alternate k-mers from k50-k96 were performed using positive strand and ambiguous stand reads as well as negative strand and ambiguous strand reads. The positive and negative strand assemblies were extended where possible, merged and then concatenated together to produce a meta-assembly contig dataset. Large scale rearrangements and gene fusions from RNA-seq libraries were identified from contigs that had high confidence GMAP (v2012-12-20) alignments to two distinct genomic regions. Evidence for the alignments were provided from aligning reads back to the contigs and from aligning reads to genomic coordinates. Events were then filtered on read thresholds. Insertions and deletions were identified by gapped alignment of contigs to the human reference using GMAP. The events were then screened against dbSNP and other variation databases to identify putative novel events.

### miRNA Sequencing

Small RNAs, containing microRNA (miRNA), in the flow-through material following mRNA purification on a MultiMACS separator (Miltenyi Biotec) are recovered by ethanol precipitation. miRNA-seq libraries are constructed using a 96-well plate-based protocol developed at the BC Cancer Agency, Genome Sciences Centre. Briefly, an adenylated single-stranded DNA 3′ adapter is selectively ligated to miRNAs using a truncated T4 RNA ligase2 (New England Biolabs). An RNA 5′ adapter is then added, using a T4 RNA ligase (Ambion) and ATP. Next, first strand cDNA is synthesized using Superscript II Reverse Transcriptase (Invitrogen), and serves as the template for PCR. Index sequences (6 nucleotides) are introduced at this PCR step to enable multiplexed pooling of miRNA libraries. PCR products are pooled, then size-selected on an in-house developed 96-channel robot to enrich the miRNA containing fraction and remove adapter contaminants. Each size-selected indexed pool is ethanol precipitated and quality checked on an Agilent Bioanalyzer DNA 1000 chip and quantified using a Qubit fluorometer (Invitrogen, cat. Q32854). Each pool is then diluted to a target concentration for cluster generation and loaded into a single lane of a HiSeq 2000 flow cell for sequencing with a 31-bp main read (for the insert) and a 7-bp read for the index.

Sequence data are separated into individual samples based on the index read sequences, and the reads undergo an initial QC assessment. Adapter sequence is then trimmed off, and the trimmed reads for each sample are aligned to the NCBI GRCh37-lite reference genome.

Routine QC assesses a subset of raw sequences from each pooled lane for the abundance of reads from each indexed sample in the pool, the proportion of reads that possibly originate from adapter dimers (i.e. a 5′ adapter joined to a 3′ adapter with no intervening biological sequence) and for the proportion of reads that map to human miRNAs. Sequencing error is estimated by a method originally developed for SAGE.

Libraries that pass this QC stage are preprocessed for alignment. While the size-selected miRNAs vary somewhat in length, typically they are ^~^21 bp long, and so are shorter than the 31-bp read length. Given this, each read sequence extends some distance into the 3’ sequencing adapter. Because this non-biological sequence can interfere with aligning the read to the reference genome, 3′ adapter sequence is identified and removed (trimmed) from a read. The adapter-trimming algorithm identifies as long an adapter sequence as possible, allowing a number of mismatches that depends on the adapter length found. A typical sequencing run yields several million reads; using only the first (5′) 15 bases of the 3′ adapter in trimming makes processing efficient, while minimizing the chance that a miRNA read will match the adapter sequence.

After each read has been processed, a summary report is generated containing the number of reads at each read length. Any trimmed read that is shorter than 15bp is discarded; remaining reads are submitted for alignment to the reference genome. BWA (Li and Durbin, 2009) alignment(s) for each read are checked with a series of three filters. A read with more than 3 alignments is discarded as too ambiguous. Only perfect alignments with no mismatches are used. Reads that fail the Illumina basecalling chastity filter are retained, while reads that have soft-clipped CIGAR strings are discarded.

For reads retained after filtering, each coordinate for each read alignment is annotated using a reference database, and requiring a minimum 3-bp overlap between the alignment and an annotation. If a read has more than one alignment location, and the annotations for these are different, we use a priority list to assign a single annotation to the read, as long as only one alignment is to a miRNA. When there are multiple alignments to different miRNAs, the read is flagged as cross-mapped (de Hoon et al., 2010), and all of its miRNA annotations are preserved, while all of its non-miRNA annotations are discarded. This ensures that all annotation information about ambiguously mapped miRNAs is retained, and allows annotation ambiguity to be addressed in downstream analyses. Note that we consider miRNAs to be cross-mapped only if they map to different miRNAs, not to functionally identical miRNAs that are expressed from different locations in the genome. Such cases are indicated by miRNA miRBase names, which can have up to 4 separate sections separated by "-", e.g. hsa-mir-26a-1. A difference in the final (e.g. ‘-1’) section denotes functionally equivalent miRNAs expressed from different regions of the genome, and we consider only the first 3 sections (e.g. ‘hsa-mir-26a’) when comparing names. As long as a read maps to multiple miRNAs for which the first 3 sections of the name are identical (e.g. hsa-mir-26a-1 and hsa-mir-26a-2), it is treated as if it maps to only one miRNA, and is not flagged as cross-mapped.

The minimum depth of sequencing required to detect the miRNAs that are expressed in one sample is 1,000,000 reads per library mapped to miRBase (v21) annotations. Finally, for each sample, the reads that correspond to particular miRNAs are summed and normalized to a million miRNA-aligned reads to generate the quantification files. TARGET and TCGA miRNA quantifications were normalized with pSVA, preserving known subtype-specific miRNA expression patterns, prior to comparison^63^.

Differentially expressed miRNAs and mRNA were determined by Wilcoxon tests, where significantly differentially expressed miRNAs were those with Benjamini-Hochberg multiple test corrected p-values <0.05. Correlation between miRNA and mRNA expression was determined using the Spearman correlation.

### DNA-methylation analysis

Bisulfite conversion of genomic DNA was performed with EZ DNA methylation Kit (Zymo Research, Irvine, CA) following the manufacturer’s protocol with modifications for the Infinium Methylation Assay. Briefly, one microgram of genomic DNA was mixed with 5 μl of Dilution Buffer and incubated at 37°C for 15 minutes and then mixed with 100 μl of conversion reagent prepared as instructed in the protocol. Mixtures were incubated in a thermocycler for 16 cycles at 95°C for 30 seconds and 50°C for 60 minutes. Bisulfite-converted DNA samples were loaded onto the provided 96-column plates for desulphonation, washing and elution. The concentration of bisulfite-converted, eluted DNA was measured by UV-absorbance using a NanoDrop-1000 (Thermo Fisher Scientific, Waltham, MA). Bisulfite-converted genomic DNA was analyzed using the Infinium Human Methylation27 Beadchip Kit (Illumina, San Diego, CA, #WG-311-1202). DNA amplification, fragmentation, array hybridization, extension and staining were performed with reagents provided in the kit according to the manufacturer’s protocol (Illumina Infinium II Methylation Assay, #WG-901-2701). Briefly, 4 μl of bisulfite-converted genomic DNA at a minimum concentration of 20 ng/μL) was added to 0.8 ml 96-well storage plate (Thermo Fisher Scientific), denatured in 0.014N sodium hydroxide, neutralized and then amplified for 20-24 hours at 37°C. Samples were fragmented at 37°C for 60 minutes and precipitated in isopropanol. Re-suspended samples were denatured in a 96-well plate heat block at 95^o^C for 20 minutes. 15 μl of each sample was loaded onto a 12-sample BeadChip, assembled in the hybridization chamber as instructed by the manufacturer and incubated at 48°C for 16-20 hours. Following hybridization, the BeadChips were washed and assembled in a fluid flow-through station for primer-extension reaction and staining with reagents and buffers provided. Polymer-coated BeadChips were scanned in an iScan scanner (Illumina) using Inf Methylation mode. For both HumanMethylation27 and HumanMethylation450 arrays, methylated and unmethylated signal intensity and detection p-values were extracted after background correction and (in the case of HumanMethylation450 arrays) dye-bias equalization by normal-exponential convolution (noob^64^) as implemented in the minfi-package^65^. Data from HumanMethylation450 arrays were additionally normalized using functional normalization (funnorm^66^) as implemented in the minfi-package, then summarized as beta values [M /(M+U)]. Probes with an annotated SNV within the CpG or single-base extension site are masked as NA across all samples. Probes with non-detection probability > 0.01 are masked as NA for individual samples.

### Transcriptional silencing evaluation and tabulation

Transcription is influenced by a large number of features, among which is methylation of genomic CpG dinucleotides, which often leads to methyl-binding domain proteins excluding transcriptional activators when it occurs near a transcription start site. Not all gene promoters are influenced by differences in DNA methylation, and not all promoters which are thusly influenced are relevant in a given cell type. Thus we sought to identify bundles of transcripts (genes) whose expression appears to be influenced by promoter CpG methylation and whose expression potential is perturbed in a subset of AML cases.

To establish a uniform criteria for “calling” such events, we evaluated over 50,000 loci from the Illumina HumanMethylation450 (“450k”) microarray near the transcription start sites of over 20,000 transcripts. Where any variance in transcript abundance was explained by variation in DNA methylation levels at a locus, we retained the locus and gene symbol for further evaluation. With this set of several thousand potential marker pairs, we iteratively sought “silencing” cutoff points, such that the maximum expression of a gene in any sample with methylation above the cutoff level was less than or equal to the median expression of samples below the cutoff. The relative levels of DNA methylation and expression appeared to differ systematically between TCGA AML and TARGET AML patients. Therefore we retained the most conservative (highest) cut-point from among the two cohorts. A large number of TARGET AML patients were previously assayed on the promoter-centric Illumina HumanMethylation27 (“27k”) microarray; to maximize the sample size for silencing calls, we performed the same conservative procedure as described above with 27k loci. Whenever a locus could be found with a suitable cut-point on both 27k and 450k arrays, we used the two loci to cross-validate transcriptional silencing behavior between the two (largely disjoint) sets of samples (TCGA AML patients were assayed on both 27k and 450k arrays, so we used the appropriate complementary assay to cross-validate each cutoff in TCGA). The resulting set of “tag CpGs” (loci with satisfactory cutoff values for a given gene) on each platform, along with the results of applying these cutoffs to dichotomize patient samples into “silenced” or not, are provided in **Table S9**. Selected loci and genes affected across multiple patients are plotted in Fig. 5a, annotated within each major cytogenetic group by the fraction of patients silenced.

### Non-negative matrix factorization, DNA methylation signature derivation, and hierarchical clustering

Non-negative matrix factorization (NMF) decomposes a strictly positive data matrix *X* (with *N* rows and *M* columns) into a lower-dimensional *NxK* weight matrix *W* and a corresponding *KxM* score matrix *H^67^*. The crux of the decomposition is to find coefficients for *W* and *H* which, when multiplied, most closely recover the original high-dimensional data matrix *X*, as there is no guarantee that a global optimum exists in the absence of further constraints. This can be approached as an optimization problem: given an estimate of the underlying rank *K* for the weight matrix *W*, what coefficients minimize the squared reconstruction error (*X*-*WH*)^2^? When this is formulated as a non-negative least squares fit, alternating between fits for *W* and *H* at each iteration, a fast sequential coordinate descent procedure implemented by Eric Xihui Lin (https://cran.r-project.org/web/packages/NNLM/vignettes/Fast-And-Versatile-NMF.html) is useful for the large matrix we use for the input *X*. To decrease the size of *X* without discarding information, the HumanMethylation450 data was further collapsed by aggregating signals at adjacent CpG sites (up to 50bp separated) using the cpgCollapse function in the minfi-package, yielding 221,406 discrete clustered methylation measurements, of which approximately half (118,586) showed non-negligible variation across diagnostic tumor samples and/or matched remission samples. The underlying identifiable rank *K* of the low-dimensional weight matrix *W* was estimated by 5-fold cross validation, using random row x column knockouts (set to NA) in 20% of the matrix entries for each fold, followed by minimization of reconstruction error and maximization of inferred rank. Based on this procedure, the optimal rank *K* (with mean absolute error of 0.02793436) for *W* was estimated as 31. By masking with *W* and *H* matrices derived from normal bone marrow populations (for which *K* was chosen as 13, again based on reconstruction error as above), we subtracted “normal” hematopoietic cell signals and simultaneously estimated the purity (cellularity) of each tumor sample, which allowed us to amplify disease-specific signals, correcting *in silico* for estimated purity on a logit scale, and finally transforming back to the original proportional 0-1 scale for presentation in Fig. 5b. The 31-row by 284-column patient score matrix *H,* are provided in **Table S10**; selected signatures of particular interest are plotted in Fig. 5b. Ward’s method was employed to cluster columns (patients) in the figure panel by Manhattan distance.

### Survival analysis

We tested an additional cohort of pediatric AML patients for outcome measures associated with alterations of *FLT3-ITD, NPM1* and *WT1* mutations and *NUP98-NSD1* translocations (Fig. 3C, lower right, and **Fig. S13**, abbreviated “DCOG”). Patient data for this cohort was provided by the Dutch Childhood Oncology group (DCOG), the AML ‘Berlin-Frankfurt-Münster’ Study Group (AML-BFM-SG), the Czech Pediatric Hematology (CPH) group, the St Louis Hospital in Paris, France, the Medical Research Council (MRC), and the Italian Association for Pediatric Hematology and Oncology (AIEOP). Patients were treated by LAME 86, DCOG/AML-BFM 87, DCOG 92-94/AML-BFM 93, AML-BFM 98, AEIOP-2002/01, ELAM02, AML-BFM 04 and MRC-12/15 protocols^68–75^. These protocols consisted of 4-5 blocks of intensive chemotherapy, using a standard cytarabine and anthracycline backbone. All patients in this cohort were previously published by Balgobind et al.^76^, and were extensively screened by RT-PCR or FISH for recurrent aberrations, such as *KMT2A*-rearrangements, *RUNX1-RUNX1T1, CBFB-MYH11, PML-RARA, NUP98*-rearrangements, *FLT3*-ITD, and mutations in *NPM1, CEBPA, WT1, N/KRAS* and *c-KIT*^76–79^, and included 326 patients with data available on *NUP98-NSD1, NPM1, FLT3-ITD* and *WT1* status. Complete remission was obtained in 74.8% of the patients. A total of 114 patients (35.0%) received a HSCT, of which 35 (10.7%) received an HSCT at first complete remission. The median follow up time of survivors was 4.5 years (range 0.3-28 years) and the cohort-wide OS and EFS were 59.5% and 41.9%, respectively.

The Kaplan-Meier method was used to estimate overall survival (OS) and event free survival (EFS). OS is defined as the time from study entry until death. EFS is defined as the time from study entry until death, induction failure, or relapse. Patients lost to follow-up were censored at their date of last known contact. Comparisons of OS and EFS were made using the log-rank test.

TARGET and TCGA subjects were combined in Cox proportional hazards fits for association of DNA methylation signatures with survival outcome, and model parameters for well-established risk factors (*TP53* mutation, white blood cell count at diagnosis) were also estimated. Due to the nonlinear association of age with survival in pediatric AML patients, and the difficulty of properly evaluating this relationship, we instead stratified the Cox proportional hazards fits by cohort.

For miRNA associated survival analyses, the expression (RPM) cut point between high and low expression groups for each miRNA was defined using the X-tile method^80^, where all separation points between patients were considered and the selected cut point was the one that provided the optimal (lowest) EFS log rank p-value.

### Life Sciences Reporting Summary

For additional information on experimental design, methods and reagents, please see the associated “Life Sciences Reporting Summary Report” file.

### Data Availability

Complete details of sample preparation protocols, clinical annotations, and all primary data are available through the TARGET Data Matrix (https://ocg.cancer.gov/programs/target/data-matrix). Sequence data are also accessible through the National Cancer Institute Genomic Data Commons ((https://portal.gdc.cancer.gov/legacy-archive/search/f) or the National Center for Biotechnology Information’s dbGaP (https://www.ncbi.nlm.nih.gov/gap) under accession number phs000218.

